# Intronic non-coding RNAs within ribosomal protein coding genes can regulate biogenesis of yeast ribosome

**DOI:** 10.1101/289751

**Authors:** Akshara Pande, Rani Sharma, Bharat Ravi Iyengar, Vinod Scaria, Beena Pillai, Samir K Brahmachari

## Abstract

The genome of the budding yeast (*Saccharomyces cerevisiae*) has selectively retained introns in ribosomal protein coding genes. The function of these introns has remained elusive in spite of experimental evidence that they are required for the fitness of yeast. Here, we computationally predict novel small RNAs that arise from the intronic regions of ribosomal protein (RP) coding genes in *Saccharomyces cerevisiae*. Further, we experimentally validated the presence of seven intronic small RNAs (isRNAs). Computational predictions suggest that these isRNAs potentially bind to the ribosomal DNA (rDNA) locus or the corresponding rRNAs. Several isRNA candidates can also interact with transcripts of transcription factors and small nucleolar RNAs (snoRNAs) involved in the regulation of rRNA expression. We propose that the isRNAs derived from intronic regions of ribosomal protein coding genes may regulate the biogenesis of the ribosome through a feed-forward loop, ensuring the coordinated regulation of the RNA and protein components of the ribosomal machinery. Ribosome biogenesis and activity are fine-tuned to the conditions in the cell by integrating nutritional signals, stress response and growth to ensure optimal fitness. The enigmatic introns of ribosomal proteins may prove to be a novel and vital link in this regulatory balancing act.

## Introduction

All the 401 introns present in its genome (Saccharomyces Genome Database (SGD)) [1] are found within less than 370 (7%) of 6000 genes in the yeast, *Saccharomyces cerevisiae*. These include introns in snoRNAs, tRNAs and the mitochondrial genome, pushing the number of spliceosomal introns to a shear 281. In contrast, the closely related *Schizosaccharomyces pombe*, has an abundance of 4698 introns in 2252 genes which constitutes about 45% of nearly 5000 genes in its genome (PomBase) [2].Compared to other eukaryotic genomes of comparable size, the *Saccharomyces cerevisiae* genome is unusually intron free. In spite of this loss of introns in the majority of yeast genes [3], 104 (out of 281) spliceosomal introns are in ribosomal protein coding genes [4].The *S. cerevisiae* genome contains 137 coordinately regulated ribosomal protein coding genes, 76% of which carry one or more introns. Evolutionary analysis of the yeast genome has shown that the Saccharomyces family (exemplified by *S. cerevisiae*) has lost introns extensively in spite of having intron rich ancestors [5]. This has led to the hypothesis that the retained introns have a functional relevance, due to which these have been evolutionarily maintained in the genome, despite the loss of introns in other gene classes[4,6].

In recent years, almost all organisms, prokaryotic or eukaryotic, have been shown to express non-coding RNAs (ncRNAs) that carry out an ever growing set of diverse functions (reviewed in [7]). In prokaryotes, some non-coding RNAs are part of innate anti-viral defense [8] while others regulate genes involved in quorum sensing and motility (reviewed in [9]). In eukaryotes, non-coding RNAs have diverse functions: microRNAs are mediators of post-transcriptional gene regulation (reviewed in [10]), while the slightly longer piRNAs are involved in protecting genomes from transposable elements (reviewed in [11]) and long-non-coding RNAs (lncRNAs) are involved in several functions ranging from chromatin organization to structural scaffolding of proteins ([12], [13]). Intronic miRNAs and miRtrons are derived during processing of messenger RNAs and participate in dynamic regulation of gene expression [14].

Introns are no longer considered as non-functional regions since it is now clear that introns may harbour functional elements [6]. This, along with the fact that a few introns are selectively retained in yeast, prompted us to explore the possibility that these intronic regions may have functional implications. We used publicly available RNA-Seq data, and found sequence reads mapping to 12 introns in the *Saccharomyces cerevisiae* genome. Seven of these show significant expression levels in at least one of the reported studies. We find that these reads, four of which are derived from introns of ribosomal protein coding genes, can arise from small RNAs that can potentially bind to the rDNA locus or rRNA transcripts. These transcripts belong to the small and large subunits of the eukaryotic ribosome or mitochondrial ribosome. Further, we could detect the expression of these small intronic RNAs (henceforth referred to as isRNAs) in wild type yeast cells. In a strain where one of the introns specifically from *IMD4* gene, was deleted, the expression of the corresponding isRNA was lost. Thus, novel non-coding RNAs that could potentially target rRNA biosynthesis or ribosome assembly are derived from introns of ribosomal protein coding genes. We propose that this constitutes a primitive mechanism of feed-forward regulation that allows for coordinated expression of ribosomal proteins and rRNA.

## Materials and Methods

RNA-Seq data analysis: To identify small RNAs, that potentially arise from intronic regions, we collected and downloaded publicly available RNA-Seq data from Short Read Archive (SRA) of *S. cerevisiae* in the European Nucleotide Archive (http://www.ebi.ac.uk/ena/). Ten RNA-Seq experiments [ENA: SRP002218 [15], ENA:SRP017603 [16], ENA:SRX111550, ENA:SRX111551, ENA:SRX111552, ENA:SRX111553, ENA:SRX391768 [17], ENA:SRX506368, ENA:SRX506456, ENA: SRX525284 [18] available from the database were used for further analysis. FastQC [19] was applied initially to check the quality of RNA-Seq data. Pre-built *S. cerevisiae* genome (version R64, 2011) package was downloaded (http://bowtie-bio.sourceforge.net/tutorial.shtml). Subsequently, SolexaQA [20] was used to filter and trim the reads of bad quality (phred score <20). Bowtie (version 0.12.9) [21] was used to align the RNA-seq data with the reference genome (*S. cerevisiae*). We designed a computational pipeline to screen all the intronic regions to identify regions, which have a read coverage of more than 10X. We downloaded the intron sequences and their positions from yeast intron databases (http://www.yeastgenome.org/, http://metarray.ucsc.edu/cgi-bin/intron/yirIntrondb and http://intron.ucsc.edu/yeast4.1/). We merged all the BAM files with read coverage >10X. Subsequently we converted BAM to BED for the specific co-ordinates and filtered out alignments that are opposite to the orientation of the gene. For instance, all + strand alignments for *RPL18A* would be removed because the gene is on the negative strand. Further, read count per nucleotide for each intron was calculated and plotted.

Potential targets of isRNA in yeast mRNA sequences were predicted using miRanda v3.3a algorithm and RNAhybrid 2.2. A ΔG cutoff of −30kcal/mol for putative small RNAs-ncRNA interaction was used. We found that putative small RNAs could bind to several regions of the RDN-locus. RiboVision software (http://apollo.chemistry.gatech.edu/RiboVision/) was used to visualize the mapping of putative small RNAs in the small subunit of rRNA and large subunit of rRNA [22]. To establish the ribosomal targets in the RDN-locus, UCSC BLAT [23] was used.

We re-shuffled isRNA sequences to generate 1000 scrambled sequences to explore the occurrence of target sites expected by chance. Potential targets of these thousand scrambled sequences were predicted using miRanda v3.3a algorithm with ΔG cutoff of −30kcal/mol. The positions of potential target sites of isRNAs checked with predicted target sites of thousand scrambled sequences with permitted overlap of 90%. The targets of isRNA from *COF1, IMD4, RPL28, RPL7B, RPL18A* and *RPS29A* (p-value < 0.1 with 90% overlap) were found to be significantly enriched compared to corresponding scrambled sequences.

Experimental Validation: Total RNA was isolated using TRIzol, followed by cDNA synthesis with the oligodT linked primer (5’-CTCAATCGTACATAGAAACAGGGATCTTTTTTTTTTTTTTTTTT-3’) in a two-step reaction using M-MuLV Reverse Transcriptase. PCR was carried out with the cDNA using gene specific forward primer (*RPL7B*-5’-TTGTTAGTATCTATTACTTG-3’; *YRA1*-5’-GACATGTTTCCCATAGCTAT-3’; *COF1*-5’TATGTCCTTTCTTTCCTCCC-3’; *RPS29A*-5’-ATCATTGCGTATGCTGGCTG-3’; *RPL18A*-5’ CTGATGTGACAAATTATTGG-3’; *RPL28*-5’-TTACGGCTATAAAAGGTAAC-3’; *IMD4*-5’-TTTCTTTTCTGGCTGGGCTG-3’) and a universal reverse primer (5’-CTCAATCGTACATAGAAACAGGGATC-3’).

## Results

We used publicly available RNA-Seq data from 10 experiments (see methods for details) to map reads to the 401 introns of *S. cerevisiae*. We selected seven putative small RNAs after filtering out candidates with less than of 10X coverage. Read depths of the seven selected cases are shown in Figure 1. In the intron of *COF1* and *RPL18A*, distinct peaks corresponding to the isRNA were seen. But in all other introns, no such peak was evident. The *IMD4* intron includes a snRNA (snR54). The abundantly expressed reads corresponding to the snRNA are clearly seen as a distinct peak within the first half of intron. At much lower levels, reads corresponding to the predicted isRNA are also seen between 260-380 nt.

**Figure 1:**
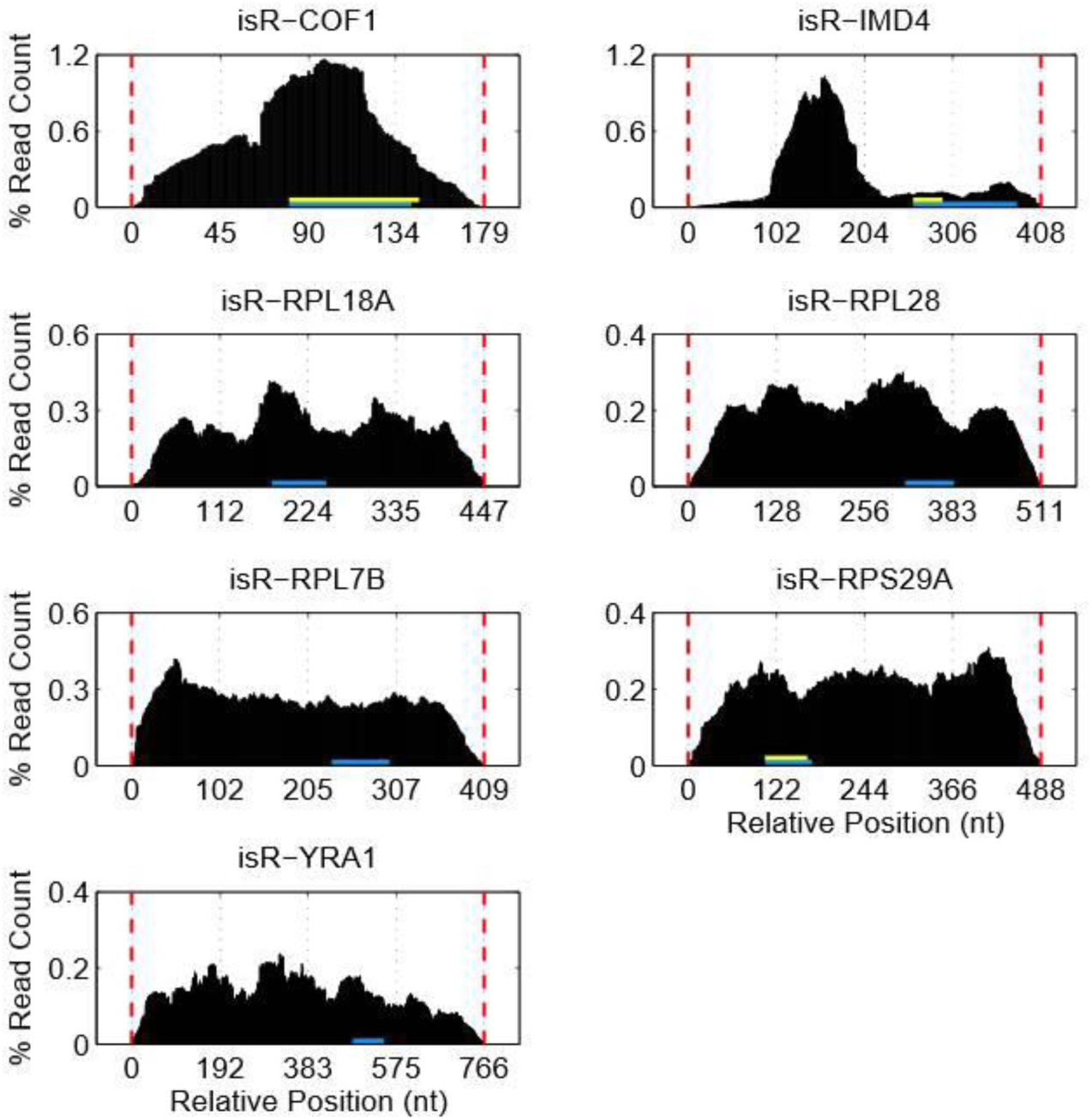
Read depth within intronic regions. X axis represents the relative position of the nucleotide from start of intron and Y axis represents the percentage of read counts. Reads from several experiments are mapped to the region between (isR-COF1) 80 to 142 nt of *COF1* gene, (isR-IMD4) 260 to 380 nt of *IMD4* gene, (isR-RPL18A) 178 to 247 nt of *RPL18A* gene, (isR-RPL28) 314 to 385 nt of *RPL28* gene, (isR-RPL7B) 232 to 299 nt of *RPL7B* gene, (isR-RPS29A) 106 to 171 nt of *RPS29A* gene and (isR-YRA1) 481 to 548 nt of *YRA1* gene. Yellow horizontal arrow shows the inferred 3’ end of isRNA and blue horizontal line shows the end of bioinformatically predicted stem loop.

If the predicted non-coding RNAs are functionally important, the region is likely to be conserved in related yeast. We used whole genome data for 11 related species (*Lachancea waltii, S. arboricola, S. bayanus, S. boulardii, S. kudriavzevii, S. mikatae, S. paradoxus, S. pastorianus*) to assess the conservation of these regions. All seven intronic non-coding RNA sequences were conserved in several Sacharomyces species, with *S. boulardii* and *S. pastorianus* showing the maximum similarity (Figure 2). We also examined the regions in the context of flanking regions to compare the degree of conservation in small RNAs with the overall level of conservation in intronic regions.

**Figure 2:**
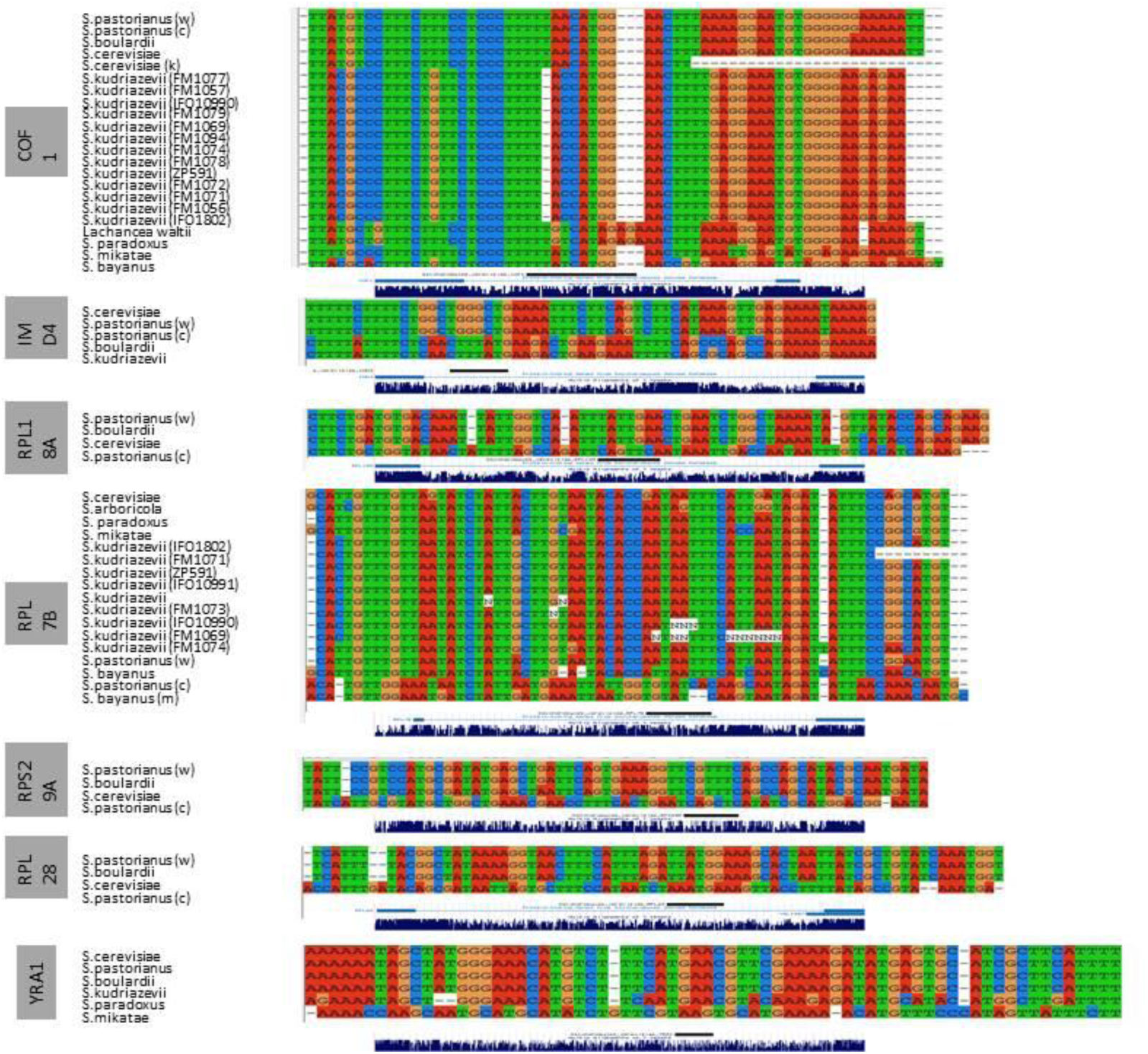
Conservation of intronic region that give rise to non-coding transcripts in *S. cerevisiae*. The sequence of the non-coding transcripts in *S. cerevisiae* was used as a query to compare to genomes of several Saccharomyces species. Multiple Sequence alignment of the results was done using ClustalX. First letter of strain name is included in parentheses wherever applicable.

We validated the presence of putative small RNAs in yeast cells, by artificially tailing it using polyA polymerase and amplifying the specific product using an isRNA specific forward primer and oligodT fused to the universal reverse primer. Using this assay (Figure 3A), we could detect the presence of seven isRNAs (*RPL7B, YRA1, COF1, RPS29A, RPL18A, RPL28* and *IMD4)*. A strain with a deletion that spans the intron of *IMD4* (originally created by J. Parenteu [4]) did not express the small RNA (Figure 3B). We also verified the presence of the isRNA derived from the intron of *RPS29A* using an RT-PCR assay with small RNA specific forward and reverse primers (Figure 3C). The observed size of the product was close to the size predicted. For instance, in the *COF1* intron (179nt), the predicted isRNA spans 82-142^nd^ position in the intron, where an increase in reads is seen (Figure 1).

**Figure 3:**
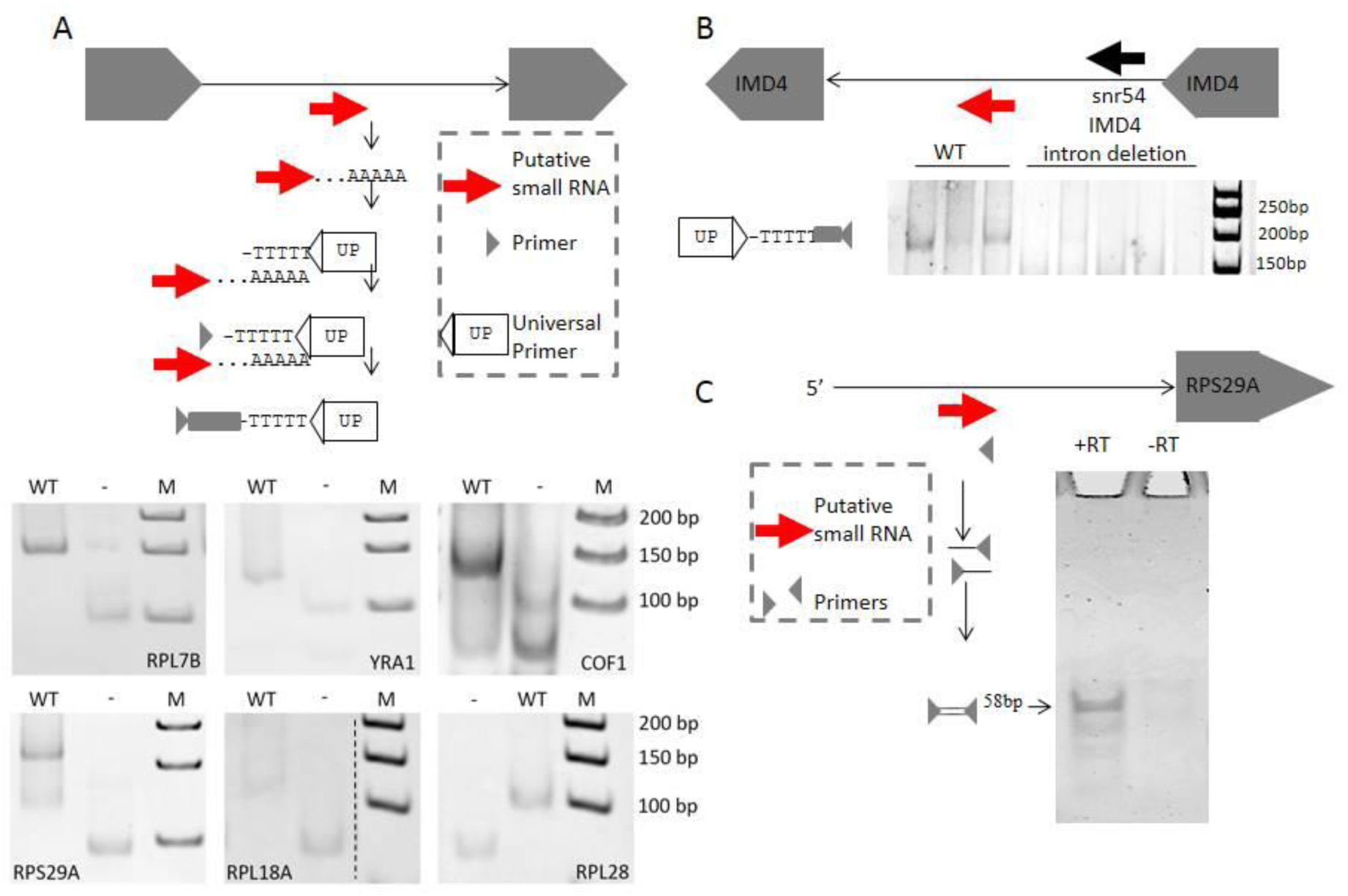
Detection of small RNA from introns in yeast. Line diagrams depict the position of the predicted small RNA (in red), and the strategy used to detect it. **(A)** In the left panel, RT PCR was carried out for the small RNA predicted from *RPL7B, YRA1, COF1, RPS29A*, RPL18A, *RPL28* and *IMD4* (+RT).DNA contamination was ruled out by repeating the experiment in the absence of reverse transcriptase (−RT). In the left panel, RNA from wild type strain was polyadenylated in vitro, and a corresponding cDNA was synthesized using a universal primer. A small RNA specific primer and the universal primer were used in combination to detect the small RNA. Control reactions were also carried out in the absence of reverse-transcriptase (−) to rule out the possibility of trace DNA contamination giving rise to the product. **(B)** In the right panel, RNA from wild type strain and a strain wherein the intron of *IMD4* was deleted were compared. The expected product was seen in three biological replicates (black line) of the wild type strain but not in the intron deleted *IMD4*.The PCR products were electrophoresed on 8% polyacrylamide gels and visualized by Ethidium bromide staining. **(C)** Intronic RNA from *RPS29A* was also detected at the predicted size using gene specific primers.

We further extended the analysis by using the same program to identify potential targets of the seven isRNAs in the non-protein coding genome. Several targets were found for each of the isRNAs (Table 2) in genomic regions that give rise to rRNA, tRNA or snoRNA. We noted that the target sites in rRNAs had a higher alignment score. The mature rRNAs are produced after post-transcriptional processing of long precursor RNAs transcribed from a single rDNA locus on chromosome 12 [24]. The locus consists of 100-200 repeating units of 9.1kb, each consisting of a 5S and a 35S precursor which get processed to form the 18S, 5.8S and 25S rRNA. Besides these rRNA coding regions, the locus consists of Internal Transcribed Spacers (ITS1, 2); External Transcribed Spacers (ETS1, 2) and Non-Transcribed Spacer regions (NTS1, 2). Therefore, we examined these 62 potential targets in the context of the rDNA locus.

**Table 1:** Details of prediction and validation of isRNAs

| isRNA (arising from intron of) | Intron size (nt) | Predicted RNA size (nt) |
| --- | --- | --- |
| COF1 | 179 | 63 |
| YRA1 | 766 | 68 |
| IMD4 | 408 | 61 |
| RPS29A | 487 | 66 |
| RPL28 | 511 | 72 |
| RPL7B | 409 | 68 |

**Table 2:** Non-coding RNAs (ncRNAs) arising from intronic region interacts with other ncRNAs

| ncRNA<br>(arising from<br>intron of) | Number of non-coding target sites<br>($\Delta G < -30$ kcal/mol) | Percent<br>of RDN<br>locus<br>covered<br>(ChrXII;<br>9.1Kb) | rRNA | tRNA | snoRNA | miscRNA<br>(snRNA,<br>lincRNA<br>etc) |
| --- | --- | --- | --- | --- | --- | --- |
| RPL28 | 129 | 5.7% | 21 | 68 | 25 | 15 |
| RPL7B | 117 | 6.8% | 27 | 31 | 42 | 17 |
| RPS29A | 99 | 4.7% | 19 | 25 | 38 | 17 |
| YRA1 | 107 | 6.0% | 21 | 8 | 57 | 21 |
| COF1 | 58 | 3.5% | 17 | - | 25 | 15 |
| RPL18A | 243 | 6.1% | 20 | 140 | 62 | 20 |
| IMD4 | 104 | 3.5% | 19 | 37 | 30 | 18 |

As shown in figure 4 the small RNAs can potentially cover a substantial region of the rDNA locus, (up to 2.64kb of the 9.1kb locus; (Supplementary File 2). To ensure that these targets may not be predicted by chance, we scrambled the isRNA sequence and predicted potential targets in the same loci. For all seven isRNA targets were found to be significantly enriched compared to corresponding scrambled sequences (p-value<0.1 with 90% overlap). The potential binding sites of the intronic small RNAs are enriched in the internal (ITS), external (ETS) and non-transcribed (NTS) spacer sequences punctuating the rDNA locus.

**Figure 4:**
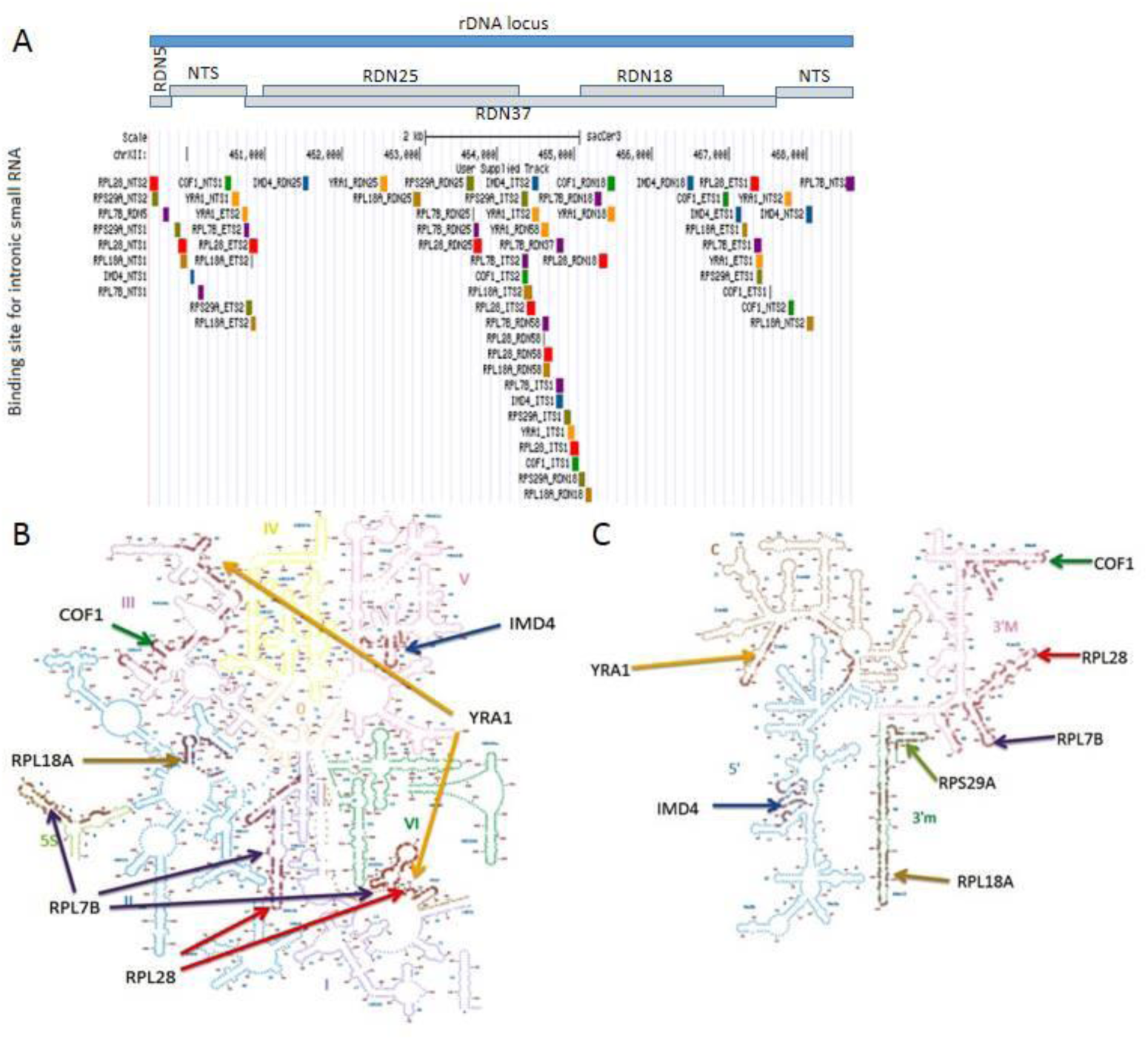
A) Potential binding sites of small RNAs in rDNA locus. Blue bars; visualized using UCSC browser. Small RNAs arising from intronic regions of different genes have been assigned specific colors (*RPL28*-red; *RPS29A*-olive; *RPL7B*-mauve, *RPL18A*-brown; *IMD4*-blue; *COF1*-green; *YRA1*-orange).For instance, the putative binding sites of the small RNA derived from *RPL28* gene (shown in red) are at NTS, ETS, ITS, RDN58, RDN18, and RDN25 regions of rDNA locus. The spacer sequences, both internal (ITS) external (ETS) and non-transcribed (NTS) punctuating the rDNA locus, are rich in putative binding sites for small RNAs. **B) Potential binding sites of small RNAs in the ribosomal RNA scaffold of the Large Subunit**: Six of seven small RNAs, labeled as per their source gene, have potential binding sites within the 5S, 5.8S and 25S subunits. **C) Potential binding sites of small RNAs in the ribosomal RNA scaffold of the Small Subunit**: All seven small RNAs, labeled as per their source gene, have potential binding sites within the 18S rRNA of the small subunit. One of the sites is potentially targeted by two overlapping small RNAs.

We visualized the putative binding sites in the context of the secondary structure of the rRNA scaffold by mapping the seven non-coding RNAs on the rRNA using the RiboVision software [22]. In the large subunit all except *RPS29A* derived ncRNA have potential binding sites unaffected by the secondary structures. The 5S rRNA is potentially targeted by isRNA from *RPL7B* while isRNAs from *YRA1, RPL28* and *RPL7B* find overlapping putative targets in the 5.8S region (Figure 4).

We searched for potential RNA-RNA hybrids between intronic small RNA and yeast mRNA based on predicted ΔG of binding [25,26]. Speculating that isRNAs may also bind to protein coding transcripts, we looked for potential targets in mRNAs of ribosome biogenesis genes. Potential targets of three isRNAs (isCOF1, isIMD4, and isYRA1) were directly related to ribosomal protein or rRNA biogenesis (Table 3). A particularly interesting target is in CURI complex involved in the transcription of ribosomal genes. Rudra et al. [27] have earlier shown that IFH1 protein shuttles between the CURI complex, which is involved in pre-rRNA processing and the RAP1 containing transcriptional complex at promoters of RP genes. This shuttling has been proposed to interface between ribosomal protein production and pre-rRNA processing. Our results suggest that the intronic RNA released during the splicing of *IMD4, YRA1* and *COF1* mRNA can potentially bind to and regulate IFH1 mRNA (Table 3).

**Table 3:**
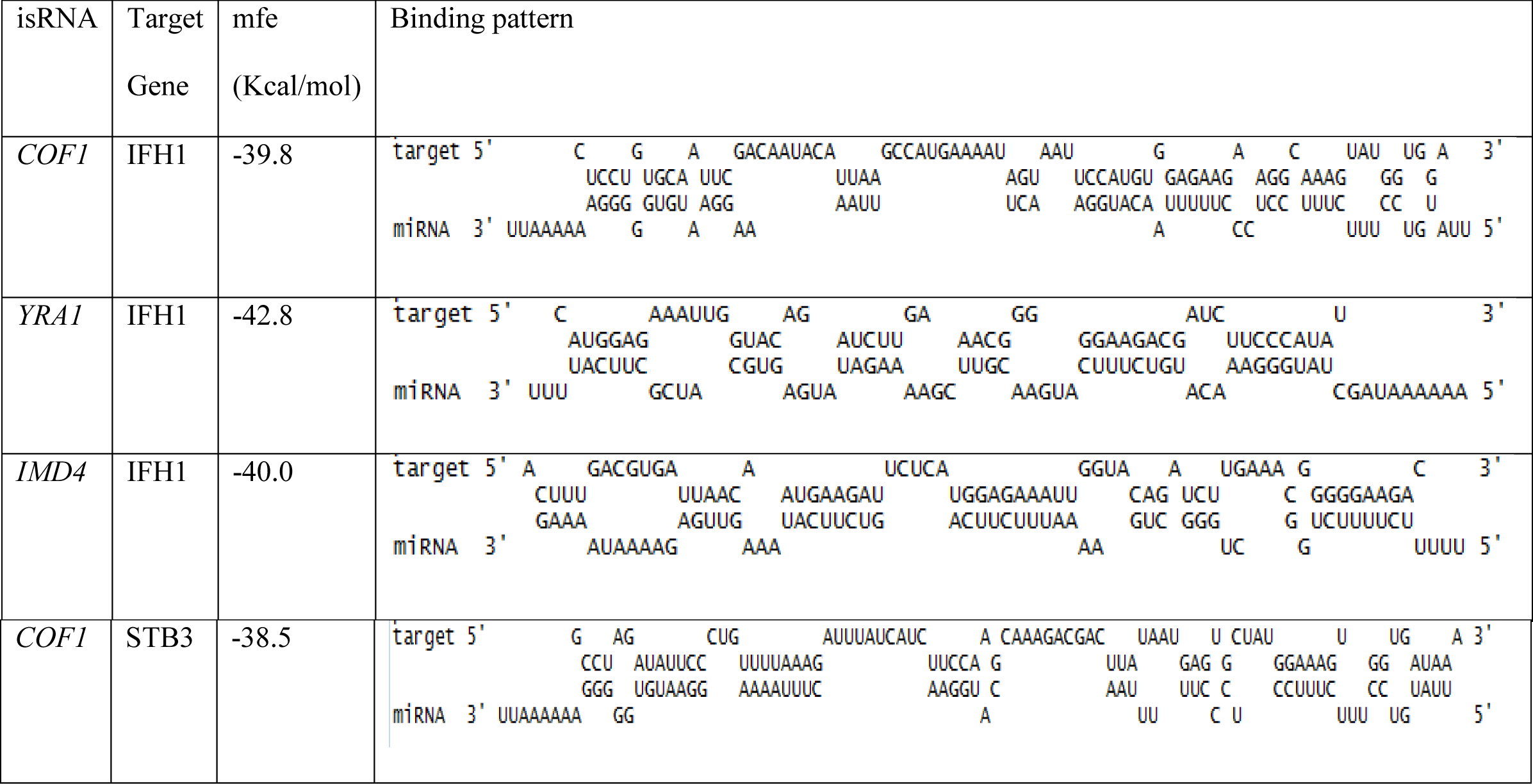
Putative interactions between non-coding RNAs and transcription factors (TF)

## Discussion

The ribosomes are amongst the most ancient and largest multi-subunit ribonucleoprotein machinery of the cell. A dedicated transcriptional machinery, run by the highly processive RNA polymerase I, produces a 6.6kb pre-rRNA transcript accounting for about 60% of the total cellular RNA transcription [28].The processing machinery then cleaves and processes this large transcript into the mature rRNA transcripts initially in the nucleus and subsequently in the cytoplasm. Folding of the rRNA, processing and assembly of ribosomal proteins are highly regulated events involving several protein factors. The production of ribosomes is closely tuned to the state of the cell by incorporating information like nutrient availability, growth, cell division and stress [29]. It is widely held that ribosomal protein production is matched to rRNA transcription and processing although the molecular basis of this crosstalk is not fully understood.

In prokaryotes, ribosomal protein biogenesis is coordinately regulated at the transcriptional level by a common promoter that regulates RP genes from a polycistronic transcript and at the translational level through a simple feedback by some of the ribosomal proteins that bind to the mRNA and prevent translation [30,31]. In eukaryotes, it is known that the RP genes and rRNA are coordinately regulated but they are neither clustered in the genome nor do they share obvious regulatory motifs. Here we show that ncRNA arising from the introns of ribosomal protein coding genes could potentially form this missing link in the coordinate regulation of ribosome biogenesis (Figure 5).

**Figure 5:**
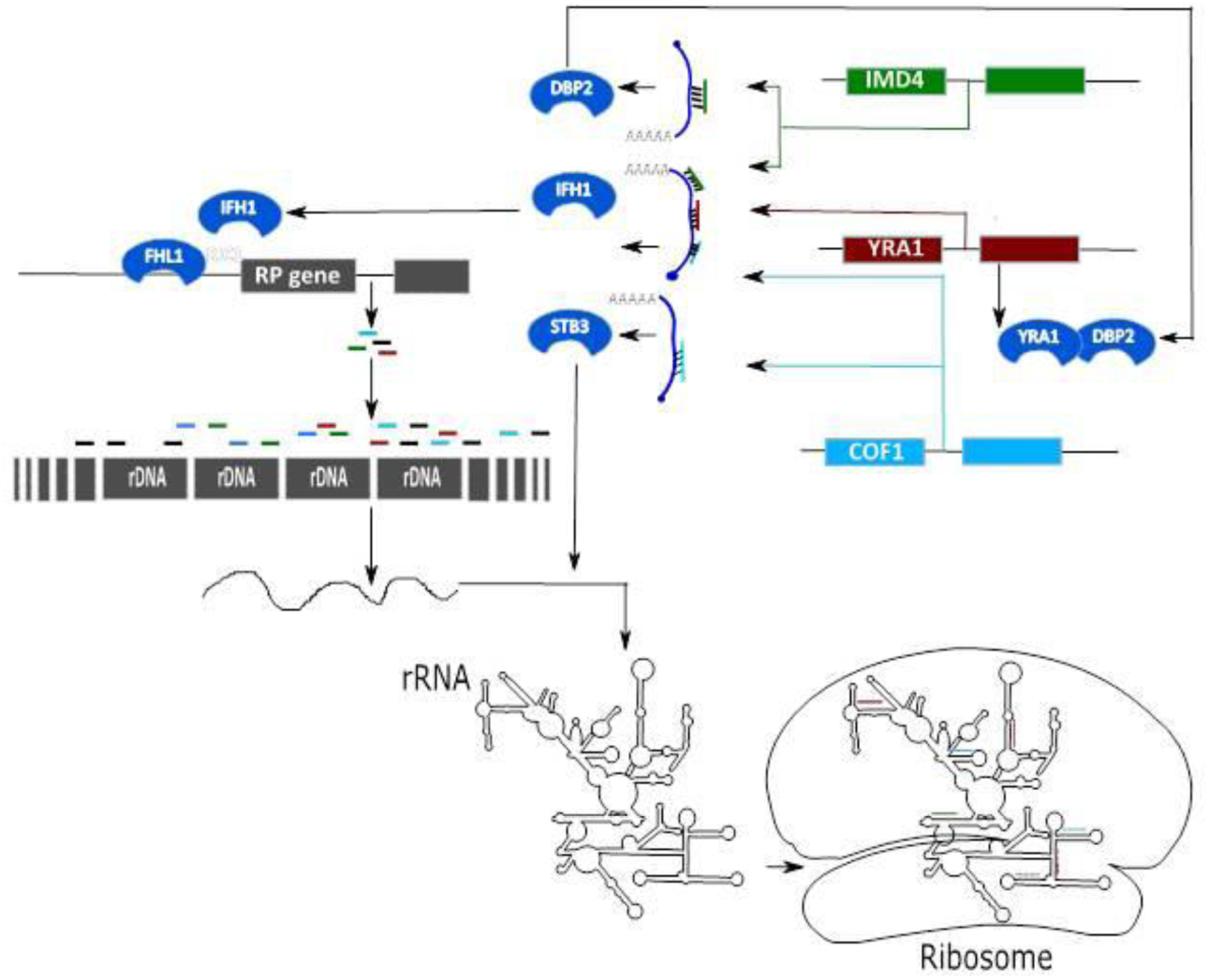
A brief overview of the hypothesized regulatory network. Small RNAs arising from intronic regions of ribosomal protein coding genes (RP genes) and *YRA1, COF1* and *IMD4* genes make a network. The small RNAs arising from introns of *YRA1, COF1* and *IMD4* can potentially target the transcription Factor IFH1, which is involved in expression of ribosomal protein genes and STB3 involved in rRNA processing. The small RNAs from RP genes have several potential binding sites in the rDNA locus and rRNA scaffold.

It was intuitively appealing to invoke the possibility that the small ncRNA could also modulate transcription factors and RNA processing enzymes that are involved in ribosome biogenesis. Such ability would allow feedback regulation of transcription and processing of the rRNA and ribosomal proteins. Three isRNAs, namely, isCOF1, isIMD4 and isYRA1 can potentially bind to the mRNA of two proteins, IFH1 and STB3 that are involved in regulation of rRNA or ribosomal protein genes. The *IFH1* mRNA has potential binding sites for all three isRNAs: isCOF1, ncIMD4 and isYRA1. The IFH1 protein is a critical component of two complexes that regulate ribosome biogenesis. Its presence in the transcriptional complex with RAP1 and Fhl1 triggers the rapid transcription of Ribosomal protein genes. When the transcription of RP genes is repressed, IFH1 leaves the promoter and localizes with the CURI complex involved in the processing of pre-rRNA [32].*STB3* codes for a protein that binds to ribosomal RNA processing element motifs and regulates gene expression [33] and its mRNA harbors potential binding sites for ncCOF1.

Interestingly, in higher mammals the first intron in some ribosomal proteins has been shown to harbour a motif [34] potentially involved in the coordinate regulation of ribosome biogenesis. Recently, Chak *et al*. [35] have shown, from deep sequencing studies, that the highly repetitive rDNA arrays of Drosophila harbor a highly conserved non-canonical miRNA. In this context, our study also implies that previously unknown non-coding RNAs may be involved in dynamic regulation of ribosome activity in *Saccharomyces cerevisiae*.

A yeast cell has about 150 tandem copies of the rDNA locus. Extrapolation of the RNA-Seq data [29] suggests that there are about 50 mRNA transcripts of each ribosomal protein per cell (3000 RP gene mRNA/60,000 mRNA transcripts: [36]), suggesting that hundreds of intronic ncRNA may be present in the cell. Furthermore, [3,6] have shown that introns are maintained in the cell even after splicing. As shown in Figure 4 the ncRNAs derived from seven introns could potentially bind to 29% of the 9.1kb rDNA locus. Thus the relative concentration of ribosomal proteins, mRNA, intronic ncRNA and rDNA locus repeats in a cell suggest that the interactions are feasible.

Hooks *et al*. 2016 found evolutionary conserved structures in YRA1, RPL28, RPL7A/B (exactly the same region as identified in this study), RPL18A/B and showed expression of the introns post splicing [6]. One measure of the relevance of these introns for biology would be the loss in fitness due to mutations. Parenteau *et al.* [4] through systematic deletion of introns in ribosomal proteins showed that the introns of ribosomal proteins are involved in expression of other ribosomal protein genes. None of the intron deletions resulted in lethality but several showed reduced fitness and altered expression of ribosomal protein genes. The deletion of the intron of *RPS29A* resulted in a significant loss of fitness [4] and an accumulation of the 20S precursor presumably due to reduced cleavage at D site between 18S and the ITS1. In our analysis, we find that the *RPS29A* derived ncRNA has a potential binding site matching this location.

The ribosome has proven to be a valuable drug target in prokaryotic systems. Many potent naturally occurring antibiotics like streptomycin, penicillin and cephalosporin act by interfering the ribosome biogenesis and function. Deletion of introns in *RPS29A*, which reduced fitness, also led to reduced sensitivity of yeast to caffeine [4]. However, high conservation of ribosomal proteins and rRNA in eukaryotes prevents its use as an anti-fungal drug target. Our findings imply that novel endogenous isRNAs may bind to the rRNA and could potentially offer valuable sites for designing small molecule inhibitors of translation.

## Abbreviations

ncRNA: non-coding RNA
isRNA: intronic small RNA
snoRNA: small nucleolar RNA
TF: transcription factor
Pre-rRNA: precursor ribosomal RNA
rDNA: ribosomal DNA
rRNA: ribosomal RNA
ITS: Internal Transcribed Spacer
ETS: External Transcribed Spacer
NTS: Non-Transcribed Spacer
nt: Nucleotide

## Authors’ Contributions

SKB designed the project. AP and BRI analyzed the data. RS carried out experimental validation. BP and SKB wrote the manuscript, VS improved the manuscript. All authors read and approved the manuscript.

## Acknowledgments

SKB acknowledges J.C. Bose National fellowship grant (GAP0093) and this work was funded by the Council of Scientific and Industrial Research (BSC0123). *IMD4* intron deletion strain was a kind gift of Julie Parenteau.

